# CITGeneDB: A comprehensive database of human and mouse genes enhancing or suppressing cold-induced thermogenesis validated by perturbation experiments in mice

**DOI:** 10.1101/250514

**Authors:** Jin Li, Su-Ping Deng, Gang Wei, Peng Yu

**Affiliations:** Department of Electrical and Computer Engineering, Texas A&M University, College Station, TX 77843, USA; TEES-AgriLife Center for Bioinformatics and Genomic Systems Engineering, Texas A&M University, College Station, TX 77843, USA; Shanghai Key Laboratory of Diabetes, Shanghai Institute for Diabetes, Shanghai Clinical Medical Centre of Diabetes, Shanghai Key Clinical Centre of Metabolic Diseases, Department of Endocrinology and Metabolism, Shanghai Jiao-Tong University Affiliated Sixth People’s Hospital, Shanghai 200233, China

## Abstract

Cold-induced thermogenesis increases energy expenditure and can reduce body weight in mammals, so the genes involved in it are thought to be potential therapeutic targets for treating obesity and diabetes. In the quest for more effective therapies, a great deal of research has been conducted to elucidate the regulatory mechanism of cold-induced thermogenesis. Over the last decade, a large number of genes that can enhance or suppress cold-induced thermogenesis have been discovered, but a comprehensive list of these genes is lacking. To fill this gap, we examined all of the annotated human and mouse genes and curated those demonstrated to enhance or suppress cold-induced thermogenesis by *in vivo* or *ex vivo* experiments in mice. The results of this highly accurate and comprehensive annotation are hosted on a database called CITGeneDB, which includes a searchable web interface to facilitate broad public use. The database will be updated as new genes are found to enhance or suppress cold-induced thermogenesis. It is expected that CITGeneDB will be a valuable resource in future explorations of the molecular mechanism of cold-induced thermogenesis, helping pave the way for new obesity and diabetes treatments. Database URL: http://citgenedb.yubiolab.org

## Introduction

Cold-induced thermogenesis (CIT) is a process by which mammals increase their resting energy expenditure in cold temperatures. CIT can be activated in two types of adipose tissues: white adipose tissue (WAT), which mainly stores fat, and brown adipose tissue (BAT), which mainly releases stored energy(1). Direct activation of BAT contributes to heat generation. In addition, induction of brown-adipocyte-like cells (beige or “brite”) in WAT depots in the process, called “browning,” also promotes heat production(2,3). Since the activation of CIT can significantly contribute to increasing resting metabolic rate in humans(4), which in turn reduces body weight, genes affecting CIT can be potential targets for antiobesity therapies.

Many genes that can affect heat production during cold exposure have been discovered. For example, Okada *et al.* found that knockout of *Acot11* increased oxygen consumption rates in both primary brown adipocytes and isolated BAT from the mutant mice, and up-regulated BAT thermogenic genes after exposure to a 4°C environment for 96 hours(5), indicating a suppressive role of *Acot11* in BAT thermogenesis. As another example, knockout of *Zfp423* decreased oxygen consumption of mouse subcutaneous WAT and down-regulated the expression of a number of WAT browning marker genes, such as *Cidea* and *Elovl3,* after exposing *Zfp423^-/-^* mice to progressively colder temperatures, suggesting that gene *Zfp423* can promote the browning of WAT under cold-exposure conditions(6). Besides these single-gene examples, some genes may function synergistically to control CIT. For example, the knockout of both *Nova1* and *Nova2* increases thermogenesis in adipose tissue upon cold exposure, but the single knockout of *Nova1* does not show a significant effect(7). These results suggest that the discovered genes may significantly contribute to uncovering the regulatory machinery of CIT.

Despite the rapid progress experienced in this field, a complete list of the genes involved in CIT is still missing. For example, the Gene Ontology (GO) Consortium does not have the term “cold-induced thermogenesis,” and the most closely related term, “adaptive thermogenesis,” (GO:1990845) has only 16 annotated mouse genes, among which the majority (13 of 16) are actually annotated to a child term, “diet induced thermogenesis.” Of the four genes *(Ucp1, Ucp2, Ucp3,* and *Pm20d1)* directly annotated to “adaptive thermogenesis,” only two *(Ucp1* and *Pm20d1)* are annotated via experimental evidence (Inferred from Mutant Phenotype (IMP)), whereas the other two are annotated via phylogenetic transfer (Inferred from Biological Ancestor (IBA)). Since *Ucp1* is involved in CIT according to the paper cited in GO for *Ucp1,* it should be annotated to “cold-induced thermogenesis” if GO had this term. These pieces of evidence confirm the incompleteness of CIT gene annotation in GO.

This lack of completeness may be explained by the fact that research in the field of CIT has been booming since 2011 **(Figure 1**), two years after BAT was identified in human adults by positron-emission tomography computed tomography (PET CT)(8), and it may be difficult for GO to keep up with the pace of progress in this specific area. Therefore, it is crucial to construct a complete list of genes enhancing/suppressing CIT, for the lack of completeness may hinder the progress of fully elucidating the mechanisms of CIT.

**Figure 1.**
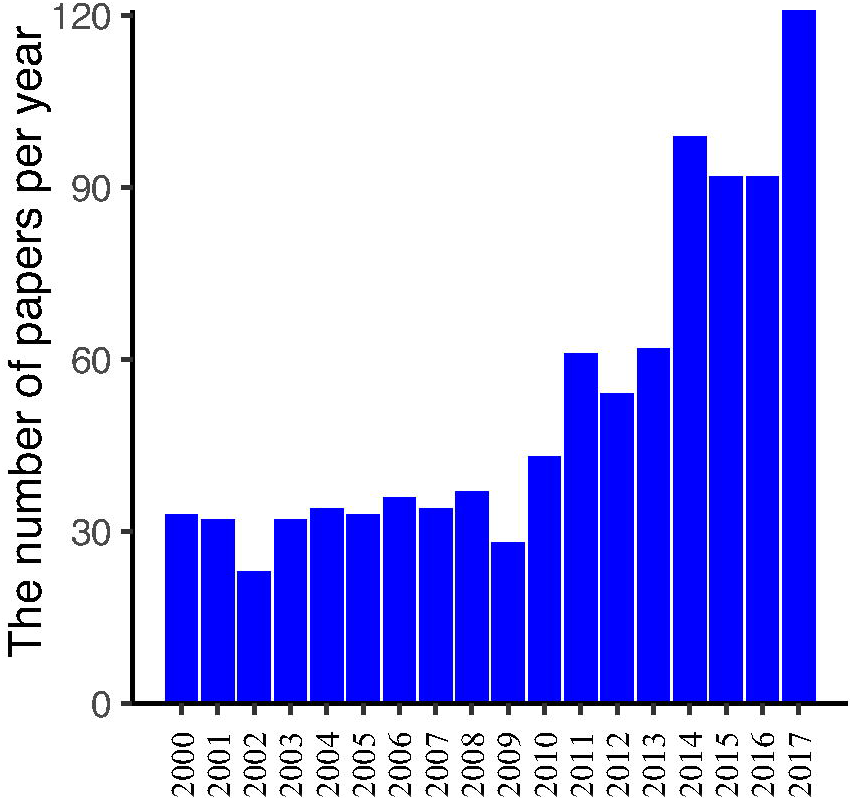
Amount of papers about “cold-induced thermogenesis” published per year since 2000. To examine the popularity of studies about thermogenesis in recent years, the number of papers was retrieved from the PubMed database by querying “cold” and “thermogenesis” in titles and abstracts on Dec. 11, 2017. The bars in the figure depict the number of papers relevant to CIT from 2000 to 2017. The number of papers published per year had been steadily relatively low before 2011, and the number increased between 2011 and 2017.

To fill this gap, we curated papers from PubMed and Google in a semiautomatic fashion. The papers show that CIT is affected when given genes are perturbed (by knockout, knockdown, overexpression, etc.). In addition, we structured the genes, the PubMed identifiers (PMIDs) of the corresponding papers, and other important data into a database called CITGeneDB.

In this paper, we introduce our effort to curate CIT-related human and mouse genes based on retrieved publications from PubMed and Google. We worked to construct and describe CIT-related data, after which we built the database CITGeneDB to host the important information of the curated CIT-related genes. In addition, we created a web interface for CITGeneDB to facilitate access to the metadata of these genes while also sharing them with various research communities.

## Data construction and data description

### Comprehensive retrieval of CIT-related papers of all the human and mouse genes validated in mice experiments

To construct a complete list of the potential papers for CIT-enhancing/suppressive human and mouse genes, all the human and mouse gene symbols were first retrieved from the HUGO Gene Nomenclature Committee (HGNC) and Mouse Genome Informatics (MGI). Each gene was searched against the PubMed database via PubMed API ESearch in Entrez Programming Utilities (E-utilities) using the query `“<gene_symbol>[tiab] AND cold[tiab] AND (thermogenic[tiab] OR thermogenesis[tiab]) NOT Review[pt] NOT Comment[pt] NOT Editorial[pt] NOT News[pt] NOT Published Erratum[pt] AND eng[la]”`. This search returned over 1,500 gene-paper pairs. Since a single paper may mention multiple genes and a gene may appear in multiple papers, we ended up with ∼200 human and mouse genes in over 1,000 papers. These results were retained for further curation for the CIT-enhancing/suppressive genes.

Some CIT-enhancing/suppressive genes may still be missing in the above search results because the titles, abstracts, and corresponding medical subject heading (MeSH) annotations used by the PubMed search for the papers describing these genes may not contain all of the keywords. To capture these missing genes, a second PubMed search for all the human and mouse genes was performed using the keywords but ignoring “cold,” which was used in the first search. This search returned ∼5,000 additional gene-paper pairs, with ∼3,000 additional papers for ∼400 additional human and mouse genes. Since these additional genes may not be related to CIT, we kept only the genes found by querying `“<gene_symbol> cold thermogenesis”` on Google. For each of these genes, Google usually does well to rank a relevant paper (if there is one) as the first hit because it uses click-through rate(9), a very effective metric for ranking webpages. Here, we checked the top three webpages for each gene kept to further ensure a high recall.

### Curation of CIT-enhancing/suppressive human and mouse genes

Our curation criterion for inclusion of a gene was that at least one thermogenesis phenotype, such as body temperature, energy expenditure, or oxygen consumption, must be significantly changed *in vivo* or *ex vivo* by the perturbation of the gene in an animal model or using tissues from an animal model in a cold-exposure condition. All animal models, such as knockout (including conditional knockout), overexpression, and drug/antibody inhibition, were considered, as long as the gene was perturbed. In other words, after the mice with a perturbed gene had been exposed to cold, some thermogenesis phenotypes were measured *in vivo* or were measured in harvesting tissues from the mice *ex vivo.* If any such phenotype was significantly changed, the perturbed gene was included. For example, the body temperature of *Hdac3* conditional knockout mice was significantly decreased in the cold condition(10), indicating that the gene can enhance heat production upon cold exposure. As another positive example, triglyceride storage of BAT harvested from *Fabp4/5* double knockout mice shrank after a 4-hour exposure to a cold environment (4°C), demonstrating that *Fabp4/5* together can promote CIT. To be stringent, genes that were tested only *in vitro* or without cold exposure were not considered. For instance, Bai *et al.* only studied *Celf1 in vitro* and not in a cold-exposure condition(11); thus, *Celf1* was not considered according to our curation criterion.

For consistency, all official gene symbols were recorded on CITGeneDB. For mice, MGI (http://www.informatics.jax.org/marker) was used to look up official symbols. Although genes hosted on CITGeneDB were mostly from mice, there are studies with human genes introduced in mice for experimentation. For these human genes, HGNC (https://www.genenames.org) was used to look up the official symbols.

To deal with special cases, we used the following approaches. When a large number of papers (e.g., >100) were returned for a gene by PubMed, the gene symbol usually was a common English word (e.g., JUN or NOV). In this case, it was not efficient to manually examine all the papers returned by PubMed, as mostly these symbols took their common English meaning in the returned papers. To overcome this limitation of PubMed, we instead searched the query term “gene <gene symbol> cold thermogenesis” on Google. Some papers could not be found by the above methods due to a lack of related keywords, but they still described genes enhancing/suppressing thermogenesis. In these cases, we manually added them to CITGeneDB.

### Statistics of enhancing/suppressive human and mouse genes in CITGeneDB

CITGeneDB is a comprehensive resource of CIT-enhancing and- suppressive human and mouse genes. Only genes confirmed in perturbation experiments using mouse models are recorded. Some information about the experiments is included in the database, such as the official symbols of the perturbed genes and perturbation type. In addition, the PMIDs of the corresponding references for each gene are stored in the database.

CITGeneDB currently has 95 CIT-enhancing genes and 47 CIT-suppressive genes. The perturbation type is knockout for most of these genes, with the exception of overexpression using adenovirus (e.g., *HOXC10,* PMID:28186086), point mutation (e.g., *Tshr,* PMID:18559984), deletion mutation (e.g., *Kdm6b,* PMID: 26625958), antibody neutralization (e.g., *Acvr2b,* PMID:22586266), and conditional transgenic overexpression (e.g., *Wnt10b,* PMID:15190075). Most genes can individually function in CIT, but some genes need to work synergistically to affect CIT. Mostly, they have redundant or similar functions, such as *Fabp4/5, Nova1/2,* and *Adrb1/2/3.* It should be noted that although our current study focuses only on CIT, the genes that were curated also may be involved in other types of thermogenesis. For example, *Ucp1* is involved in diet-induced thermogenesis. Moreover, CITGeneDB will be periodically updated when more human or mouse genes are found in new literature. Continuously updating this database will likely maintain its impact on obesity and diabetes research.

### Web interface for CITGeneDB

To facilitate database access, we developed a web interface that allows users to browse and search. Users can select the number of entries (label 1) to show on a page (**Figure 2**). For each entry, the main information (label 2) includes “Official symbol” (from MGI or HGNC), “PubMed IDs” (of the papers supporting the thermogenesis role of the genes), “Effect” (whether the gene enhances or suppresses thermogenesis), “Genotype” (what genes were perturbed and how the genes were perturbed), and Phenotype (affected phenotypes supported by experiments). Each column can be sorted alphabetically by clicking the corresponding information bars, and keywords can be searched in all the columns of the table via the search box (label 3) to obtain the corresponding entries. For example, the result entry is shown in **Figure 3** for the search “Fabp4-cre enhancing.” Nine entries were returned that have been experimentally tested as enhancing roles in thermogenesis upon cold exposure using *Fabp4*-cre-based conditional knockout mice.

**Figure 2.**
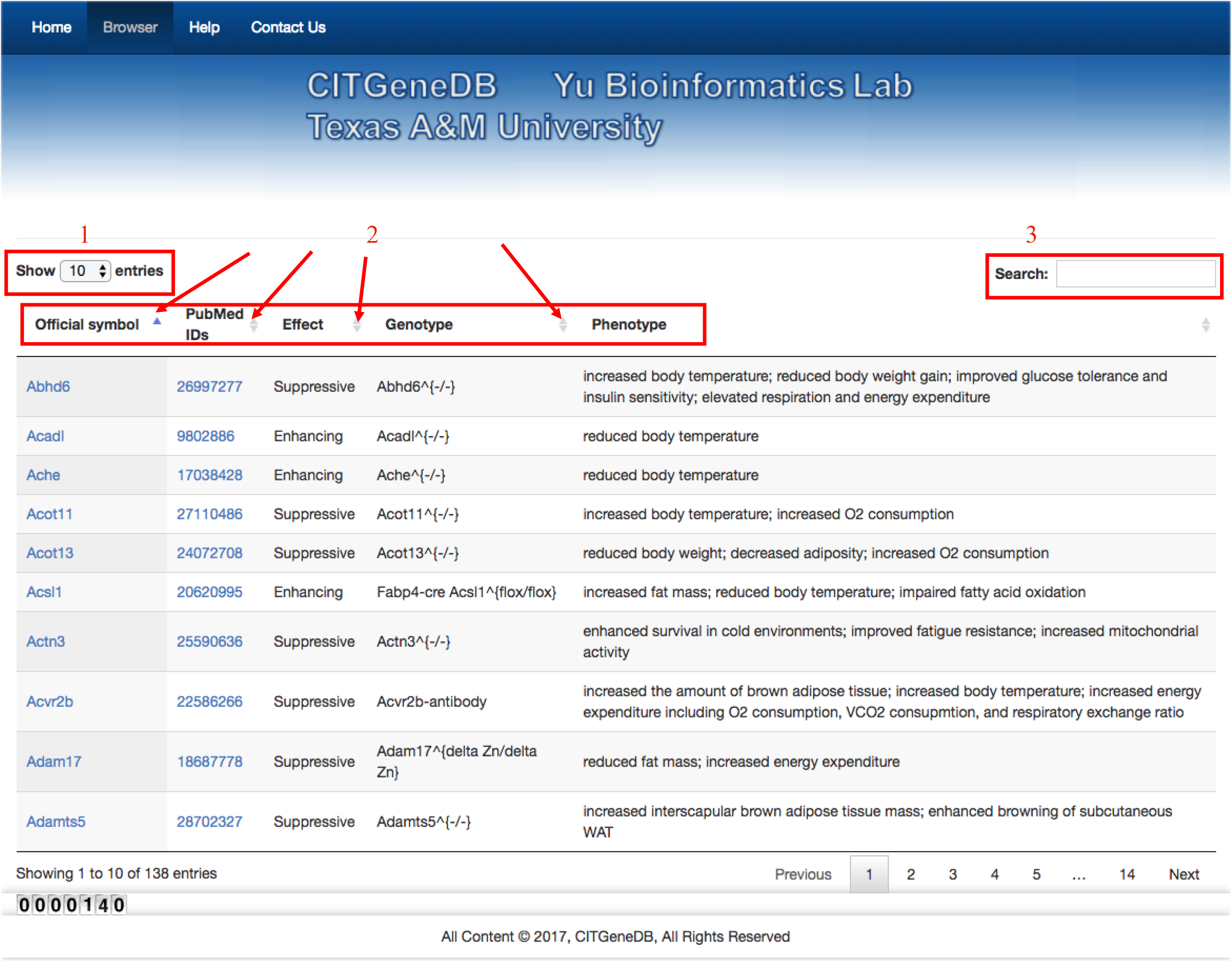
Web interface of CITGeneDB. To share the CIT-enhancing/suppressive genes, the CITGeneDB web interface was created. In the figure, label 1 is for setting the maximum number of entries on one page. Label 2 represents the main information of CIT genes including official symbols (the official gene symbol from MGI or HGNC), PMIDs of the papers supporting the thermogenesis role of the genes, Effect (whether the gene enhances or suppresses thermogenesis), Genotype (what genes were perturbed and how the genes were perturbed), and Phenotype (affected phenotypes supported by experiments). Label 3 provides the search box for the inquiry about CIT-enhancing/suppressive genes.

**Figure 3.**
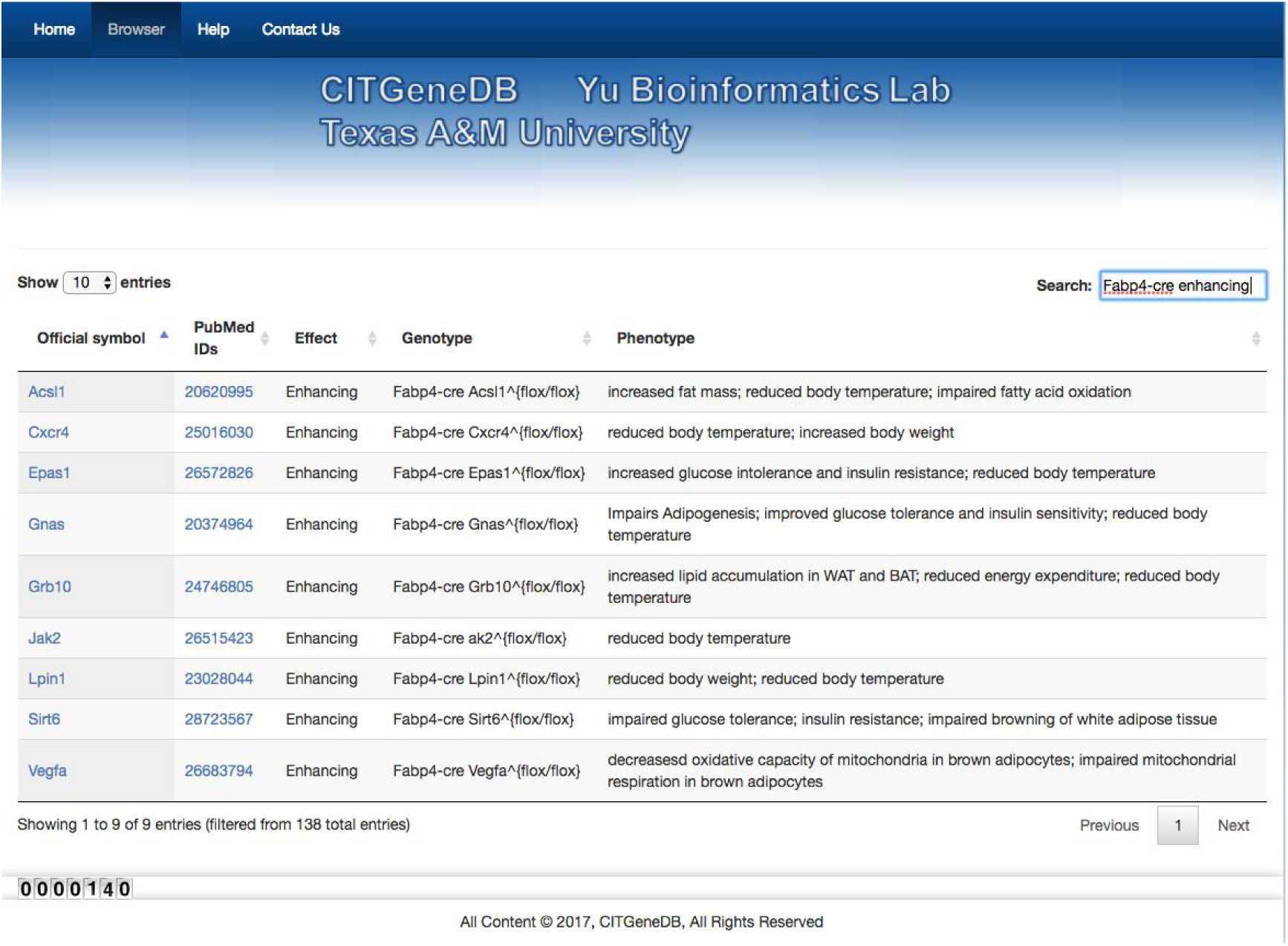
Search result example. When the keyword “Fabp4-cre enhancing” was searched, nine entries were returned. *Acsll, Cxcr4, Epasl, Gnas, Grb10, Jak2, Lpinl, Sirt6,* and *Vegfa* were all demonstrated to enhance thermogenesis in cold conditions using the fabp4-cre-based conditional knockout mouse models.

## Conclusions

CITGeneDB is a highly accurate database for CIT-enhancing/suppressive human and mouse genes derived from the literature. It is the first comprehensive resource for this purpose and can greatly complement the incomplete annotation in the related GO term. It can serve as a benchmark dataset for evaluating computational predictions of CIT-enhancing/suppressive genes and can facilitate the study of CIT from a systems biology perspective. Thus, CITGeneDB will be a valuable resource for a greater understanding of CIT and is expected to help further elucidate the regulatory networks of CIT.

## Acknowledgements

The authors thank Dr. Srujana Rayalam for her input to the data curation result.

## Funding

This work was supported by startup funding to P.Y. from the ECE department and Texas A&M Engineering Experiment Station/Dwight Look College of Engineering at Texas A&M University and by funding from TEES-AgriLife Center for Bioinformatics and Genomic Systems Engineering (CBGSE) at Texas A&M University, by TEES seed grant, and by Texas A&M University-CAPES Research Grant Program.

*Conflict of Interest:* none declared.

## References

1. Ellis, J.M., Li, L.O., Wu, P.C., et al. (2010) Adipose acyl-CoA synthetase-1 directs fatty acids toward beta-oxidation and is required for cold thermogenesis. Cell metabolism, 12, 53–64.

2. Berry, R., Rodeheffer, M.S. (2013) Characterization of the adipocyte cellular lineage in vivo. Nat Cell Biol, 15, 302–308.

3. Seale, P., Bjork, B., Yang, W., et al. (2008) PRDM16 controls a brown fat/skeletal muscle switch. Nature, 454, 961–967.

4. Ouellet, V., Labbe, S.M., Blondin, D.P., et al. (2012) Brown adipose tissue oxidative metabolism contributes to energy expenditure during acute cold exposure in humans. The Journal of clinical investigation, 122, 545–552.

5. Okada, K., LeClair, K.B., Zhang, Y., et al. (2016) Thioesterase superfamily member 1 suppresses cold thermogenesis by limiting the oxidation of lipid droplet-derived fatty acids in brown adipose tissue. Mol Metab, 5, 340–351.

6. Shao, M.L., Ishibashi, J., Kusminski, C.M., et al. (2016) Zfp423 Maintains White Adipocyte Identity through Suppression of the Beige Cell Thermogenic Gene Program. Cell metabolism, 23, 1167–1184.

7. Vernia, S., Edwards, Y.J., Han, M.S., et al. (2016) An alternative splicing program promotes adipose tissue thermogenesis. Elife, 5.

8. Cypess, A.M., Lehman, S., Williams, G., et al. (2009) Identification and importance of brown adipose tissue in adult humans. N Engl J Med, 360, 1509–1517.

9. Zhao, H.W., Huang, Y.F. (2012) An improved method for combination feature selection in web click-through data mining. Int Conf Inform Sci, 381–385.

10. Emmett, M.J., Lim, H.W., Jager, J., et al. (2017) Histone deacetylase 3 prepares brown adipose tissue for acute thermogenic challenge. Nature, 546, 544–548.

11. Bai, Z., Chai, X.R., Yoon, M.J., et al. (2017) Dynamic transcriptome changes during adipose tissue energy expenditure reveal critical roles for long noncoding RNA regulators. PLoS biology, 15, e2002176.

